# WRKY33-mediated indole glucosinolate metabolic pathway confers host resistance against *Alternaria brassicicola*

**DOI:** 10.1101/2021.04.22.440953

**Authors:** Han Tao, Huiying Miao, Lili Chen, Mengyu Wang, Chuchu Xia, Wei Zeng, Yubo Li, Shuqun Zhang, Chuanyou Li, Qiaomei Wang

## Abstract

The tryptophan (Trp)-derived plant secondary metabolites, including camalexin, 4-hydroxy-indole-3-carbonylnitrile (4OH-ICN), and indole glucosinolate (IGS), show broad-spectrum antifungal activity. However, the upstream regulators of these metabolic pathways among different plant species in response to fungus infection are rarely studied. In this study, our results revealed a positive role of WRKY33 in host resistance to *Alternaria brassicicola* by directly regulating the transcription of genes involved in the biosynthesis and atypical hydrolysis of IGS both in *Arabidopsis* and Chinese kale. Indole-3-yl-methylglucosinolate (I3G) and 4-methoxyindole-3-yl-methylglucosinolate (4MI3G) are the main components of IGS. WRKY33 induces the expression of *MYB51* and *CYP83B1* which promotes the biosynthesis of I3G, the precursor of 4MI3G. Moreover, it also directly activates the expression of *CYP81F2*, *IGMT1*, and *IGMT2* to drive side chain modification of I3G to produce 4MI3G, which is in turn hydrolyzed by *PEN2.* However, Chinese kale showed a more severe symptom than *Arabidopsis* when infected by *Alternaria brassicicola*. Comparative analyses of the origin and evolution of Trp-metabolism indicate that the loss of camalexin biosynthesis in Brassica crops during evolution might attenuate the resistance of crops to *Alternaria brassicicola*. As a result, IGS metabolic pathway mediated by WRKY33 becomes essential for Chinese kale to deter *Alternaria brassicicola*. Our results highlight the differential regulation of Trp-derived camalexin and IGS biosynthetic pathways in plant immunity between *Arabidopsis* and Brassica crops.

**One-sentence Summary:** Pathogen-responsive WRKY33 directly regulates indole glucosinolates biosynthesis and atypical hydrolysis, conferring to host resistance to *Alternaria brassicicola* in *Arabidopsis* and Brassica crops.

## Introduction

Three pathogen-inducible tryptophan (Trp)-derived secondary metabolites, including camalexin, 4-hydroxy-indole-3-carbonylnitrile (4OH-ICN), and indolic glucosinolate (IGS), are important components of the innate immune system in model plant *Arabidopsis* (Bednarek et al., 2009; Clay et al., 2009; Rajniak et al., 2015; Pastorczyk et al., 2019). However, most Brassica crops, including *Brassica rapa* and *Brassica oleracea*, are lack of 4OH-ICN and camalexin, but rich in glucosinolates (Barco et al., 2019). In most Brassica plants, IGS metabolism is evolutionarily ancient and has been largely retained (Bednarek et al., 2011; Edger et al., 2015). Glucosinolate metabolic pathway is crucial for the innate immune system and their transcriptional regulatory network have been studied (Gigolashvili et al., 2007; Gigolashvili et al., 2007; Frerigmann and Gigolashvili, 2014; Li et al., 2014; Li et al., 2020). As a diverse family of secondary metabolites, glucosinolates can be divided into three groups, indolic, aliphatic, and aromatic, depending on their amino acid precursors (Grubb and Abel, 2006; Sønderby et al., 2010). They can be hydrolyzed either by typical myrosinases (TGGs) upon tissue damage, or atypical myrosinases (PENETRATION2, PEN2) under pathogen attack in living cells (Grubb and Abel, 2006; Sønderby et al., 2010). Glucosinolates and their breakdown products are critical in plant defense against pathogens (Bednarek et al., 2009; Clay et al., 2009). PEN2-dependent breakdown product of 4-methoxyindole-3-yl-methylglucosinolate (4MI3G) has been reported to function in plant defense against various fungal and oomycete pathogens (Bednarek et al., 2009).

Production of plant defense chemicals is dependent on a regulatory network that responds to pathogen invasion by activating defense-responsive transcription factors (TFs). WRKY TFs have been shown to play a vital role in modulating the cellular responses triggered by pathogens in many plant species (Dong et al., 2003; Eulgem and Somssich, 2007; Chi et al., 2013). *WRKY18*, *WRKY33*, and *WRKY40* were among the most strongly induced WRKY family genes after flg22 treatment in *Arabidopsis* (Cyril Zipfel, 2004), and these three WRKYs have been widely investigated for their roles in plant resistance to diverse pathogens (Chen et al., 2010; Birkenbihl et al., 2017). WRKY33 is a member of the WRKY superfamily, characterized by the heptapeptide WRKYGQK and zinc-finger motif. Previous researches have shown that WRKY33 regulates plant hormone signal transduction and Trp-derived secondary metabolites (4OH-ICN and camalexin) during pathogen infection (Mao et al., 2012; Barco et al., 2019).

*Alternaria brassicicola* (*A. brassicicola*), a necrotrophic pathogen, causes the black spot disease of Brassica plants which is also routinely used as a model necrotrophic pathogen (Nowicki et al., 2012). *A. brassicicola* actively kills host tissue as it colonizes and thrives on the contents of dead cells. In particular, it is known to produce host-specific and/or non-host-specific toxic toxins (Friesen et al., 2008). The phytoalexins such as brassinin and camalexin contribute to protect Brassica plants against *A. brassicicola* (Pedras & Minic, 2012). Glucosinolate-derived isothiocyanates could cause cell death in *A. brassicicola* (Nowicki et al., 2012). To date, agricultural management practices with chemical control are the main methods for controlling Alternaria black spot in Brassica crops, which are less effective and cause pollution to the environment (Martos et al., 2016). Better understanding of innate resistance mechanism of Brassica plants to this devastating pathogen is urgently needed to develop green and sustainable disease control strategies.

In comparison with the extensive studies in model plant *Arabidopsis*, the mechanisms of host resistance to *A. brassicicola* in Brassica crops remain to be further investigated. In this paper, we found that Chinese kale, an original Chinese vegetable showed more severe symptoms than *Arabidopsis* in response to *A. brassicicola.* We conducted RNA-seq analysis to identify genes responsive to *A. brassicicola* infection in *Arabidopsis* genome. Trp-derived secondary metabolic pathway genes and WRKY33 were strongly induced by *A. brassicicola*. We then further studied their functions in the resistance of Brassica plants to this fungus. Genetic and molecular studies revealed that WRKY33 regulates IGS biosynthesis and PEN2-mediated IGS metabolism both in *Arabidopsis* and Chinese kale to trigger the innate immunity in response to *A. brassicicola.* Our study also highlights WRKY33 acts as a master regulator to promote the biosynthesis of three Trp-derived defense-related metabolites including camalexin, 4OH-ICN and IGS in *Arabidopsis*, but only IGS in Brassica crops.

## Results

### Genes involved in tryptophan-derived secondary metabolic pathways and WRKY TFs are induced in *Arabidopsis* in response to *Alternaria brassicicola*

To investigate the resistance mechanism of Brassica plants to *Alternaria brassicicola* (*A. brassicicola*), we conducted disease symptom assays on the model plant *Arabidopsis* (Col-0) and Chinese kale (*Brassica oleracea* var. *alboglabra* Bailey). We found that Chinese kale showed more severe symptoms (Figure 1A), and the fungus produced approximately threefold more spores on Chinese kale plants than on *Arabidopsis* (Figure 1B), suggesting the existence of differential mechanisms in host resistance between *Arabidopsis* and Brassica crops in response to *A. brassicicola*.

**Figure 1.**
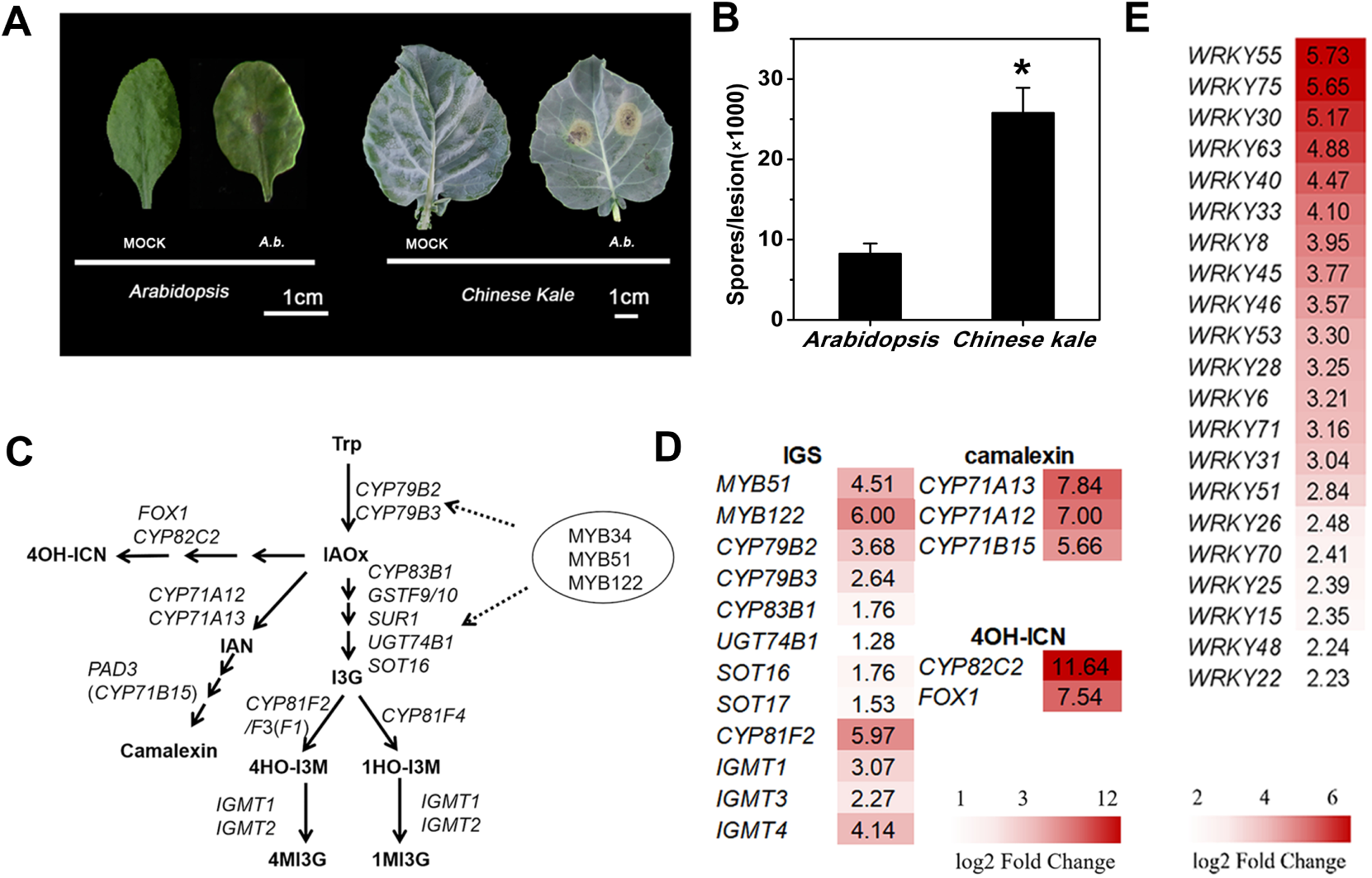
Tryptophan-derived secondary metabolic pathway genes and WRKY TFs are induced in host resistance against *Alternaria brassicicola*. **A, B:** Disease symptoms (A) and mean number of spores (B) in *Arabidopsis* and Chinese kale at 5 days after infiltration with *A. brassicicola* (*A.b.*). The asterisk indicates significant differences (p < 0.05). **C:** Simplified scheme of Trp-derived secondary metabolites pathways and the related genes in *Arabidopsis*. **D, E:** Induction of Trp-derived secondary metabolic pathway genes (D) and WRKY TFs (E) in 10 days old wild-type (Col-0) at 24 h after infiltration with *A. brassicicola* identified by RNA-seq.

RNA-seq analysis was then conducted on *Arabidopsis* (Col-0) plants with *A. brassicicola* infection to identify potential pathways and genes involved in host resistance to *A. brassicicola*. Two-fold or more (p≤0.05) change in expression of 4163 genes was observed after *A. brassicicola* infection in comparison with mock, with 2326 genes being up-regulated and 1837 genes being down-regulated (Table. S1). Tryptophan (Trp)-derived secondary metabolic pathway genes were observed during *A. brassicicola* infection (Figure 1C and 1D). The expression of *PAD3* (*CYP71B15*), encoding a P450 enzyme that catalyzes the last step of camalexin biosynthesis (Zhou et al., 1999), was strongly induced. Similarly, the mRNA level of *CYP82C2*, encoding the enzyme which converts indole-3-cyanohydrin (ICY) to 4OH-ICN, as well as a number of indolic glucosinolate (IGS) biosynthetic genes were also strongly induced (Rajniak et al., 2015). Expression of genes involved in several different transcription factors (TF) families, such as WRKYs, bHLHs, ERFs and NACs, are altered in response to *A. brassicicola* infection as well (Table. S1). We also observed that 21 WRKY TF genes were strongly (>4-fold) induced by *A. brassicicola* treatment (Figure 1E).

### WRKY33 positively regulates host resistance to *Alternaria brassicicola* both in *Arabidopsis* and Chinese kale

Among the WRKY TFs, WRKY33 has been reported to function in resistance of *Arabidopsis* to necrotrophic fungus such as *Botrytis cinerea* (Zheng et al., 2006; Birkenbihl et al., 2012). WRKY33 was induced more than 16-fold based on our RNA-seq data (Figure 1E), which suggested that *WRKY33* might function in host resistance to *A. brassicicola.* We then investigated the role of WRKY33 in host defense against *A. brassicicola* infection in *Arabidopsis*. Firstly, we carried out a time-course expression analysis of *WRKY33* in wild-type *Arabidopsis* infected by *A. brassicicola*. The expression of *WRKY33* was induced by *A. brassicicola,* and peaked at 12 h (Figure 2A). Secondly, we tried to assess the contribution of WRKY33 in *Arabidopsis* resistance to *A. brassicicola*, and performed disease assay using *wrky33* mutants. Lesions in *wrky33* mutants were almost three times as large as that in WT, which indicated that *wrky33* mutants were more susceptible than WT in response to *A. brassicicola* infection (Figure 2A). Finally, to determine whether the symptom was linked to the mutation, we expressed an AtWRKY33-YFP fusion protein under the control of the 35S promoter in the *wrky33* mutant. The phenotype of mutant was restored, as indicated by increased resistance to *A. brassicicola* than *wrky33* (Figure 2A), confirming that mutation of the *WRKY33* resulted in the phenotype of *wrky33*.

**Figure 2.**
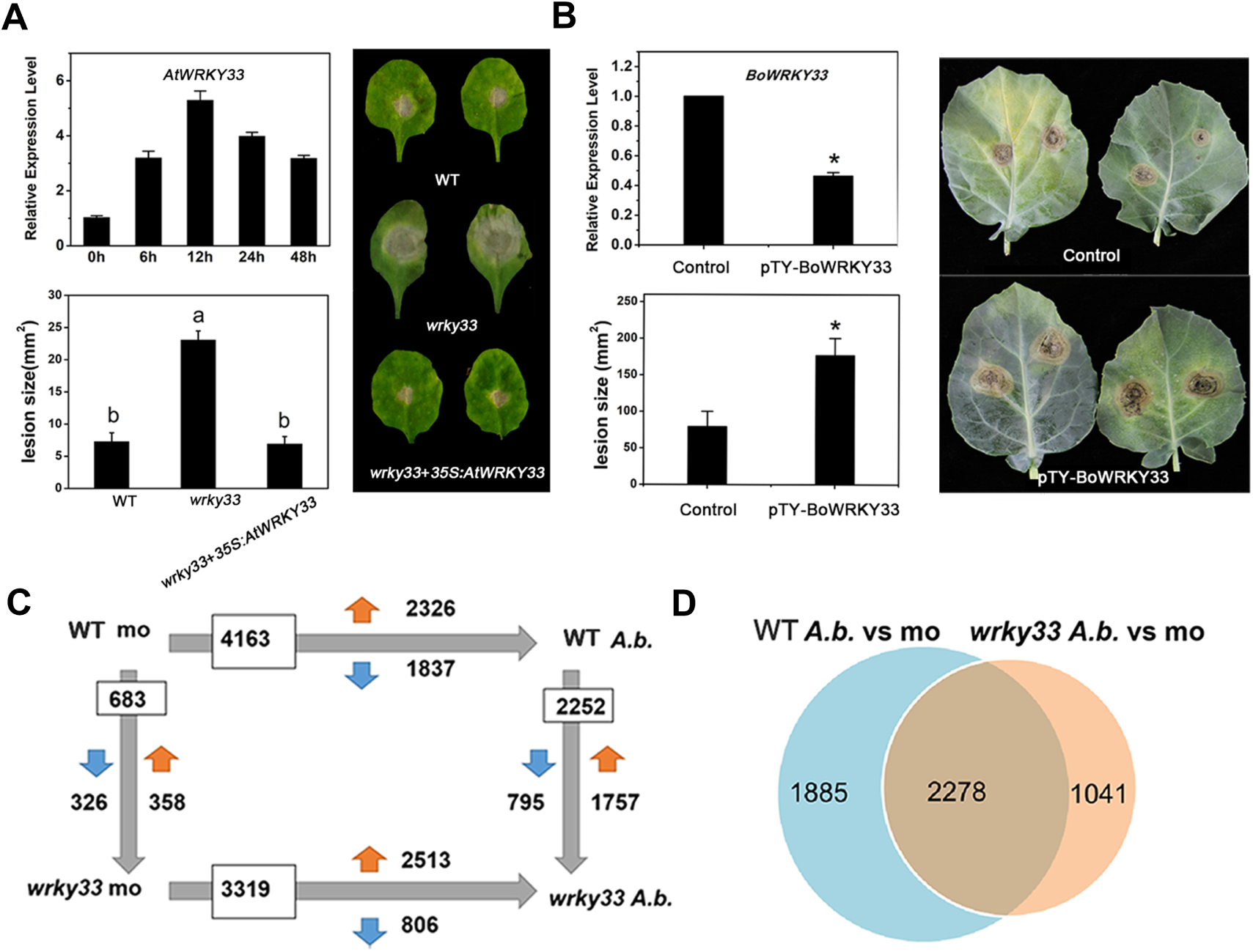
WRKY33 positively regulates host resistance in response to *Alternaria brassicicola*. **A:** qRT-PCR analysis of *WRKY33* in wild type (WT) and disease symptoms of inoculated plants of 4 weeks WT, *wrky33* and *wrky33*+*35S:AtWRKY33*. **B:** qRT-PCR analysis of *BoWRKY33* and disease symptoms of control plant with empty pTY vector (Control) and *BoWRKY33*-silenced (pTY-BoWRKY33) Chinese kale plants inoculation with *A. brassicicola*. Data for the pTY-BoWRKY33 Chinese kale are representative of three independent transgenic lines. **C:** Number of differentially expressed genes (FC≥ two fold; p ≤ 0.05) between 10 days old WT and *wrky33* at 24 h after mock treatment (mo) or infiltration with *A. brassicicola* (*A.b.*) identified by RNA-seq. Indicated are total numbers (boxed) and numbers of up-regulated (↑) and down-regulated (↓) genes between treatments or genotypes. **D:** Venn diagram illustrating the total numbers and the number of common genes affected in 10 days old WT and *wrky33* 24 h post *A. brassicicola* inoculation.

To verify whether WRKY33 also functions in resistance of Brassica crops to *A. brassicicola,* we conducted disease symptom assays in Chinese kale. The virus-induced gene silencing (VIGS) system was employed to generate Chinese kale materials with more than 50% decreased expression of *BoWRKY33* (Figure 2B). Next, we inoculated the leaves of *BoWRKY33-*silenced (pTY-BoWRKY33) and control plants with *A. brassicicola* to quantify the disease resistance, and recorded the lesion size. *BoWRKY33-*silenced plants showed more severe symptoms with lesions that were about 3 times larger than that in the control (Figure 2B). Thus, these results suggest that WRKY33 plays an important role in host defense against *A. brassicicola* infection.

To identify genes involved in WRKY33-dependent host response to *A. brassicicola* on a genome-wide level, we again performed RNA-seq analysis and examined WRKY33-mediated gene expression changes in mock, along with *A. brassicicola* treated *wrky33* and WT plants. A total of 2252 WRKY33-dependent differentially expressed genes (DEG) were identified between *A. brassicicola* infected *wrky33* and WT plants, with 1757 being up-regulated, and 795 being down-regulated in *wrky33* mutants (Figure 2C). A common set of 2278 genes showed changes in transcript level after inoculation with *A. brassicicola* in both WT and *wrky33*. The expression of 1885 genes was altered only in WT upon infection, while differential expression of 1041 genes was caused by mutation of *WRKY33* (Figure 2D). To investigate the relevance of WRKY33 in plant defense responses, we analyzed Gene Ontology (GO) term enrichment and found that WRKY33-regulated genes were significantly enriched in biological processes (BP) and molecular functions (MF) related to secondary metabolic process, response to stimuli, heme binding, and oxidoreductase activity, and many of these genes are repressed in *wrky33* mutants upon *A. brassicicola* infection (Supplemental Figure 1), suggesting that WRKY33 might enhance the host resistance to *A. brassicicola* infection via regulation of the secondary metabolic pathway.

### Indolic glucosinolate pathway is important in host resistance to *Alternaria brassicicola* in *Arabidopsis*

Trp-derived secondary metabolites (camalexin, 4OH-ICN and IGS) are important components of plant innate immune system, and WRKY33 was reported as an important regulator of camalexin and 4OH-ICN (Mao et al., 2012; Barco et al., 2019; Barco and Clay, 2020) (Figure 3A). To investigate the role of different Trp-derived metabolites in host resistance to *A. brassicicola,* the temporal expression patterns of Trp-derived secondary metabolic pathway genes were analyzed by qRT-PCR analysis with samples collected at different time points after inoculation with *A. brassicicola*. Expression of genes involved in camalexin (*CYP71A12, CYP71A13*, and *PAD3*) and 4-OH-ICN (*CYP82C2* and *FOX1*) biosynthesis was increased drastically during the infection (Supplemental Figure 2). MYB34, MYB51, and MYB122 are three transcription factors that regulate biosynthesis of indolic glucosinolates, with MYB51 being the most crucial one (Frerigmann and Gigolashvili, 2014). The mRNA levels of *MYB 51* and *MYB 122* were increased upon infection (Figure 3B). However, there was no dramatic change in expression level of *MYB34* being recorded (Figure 3B). The cytochrome P450 enzymes, CYP79B2 and CYP79B3, convert Trp to indole-3-acetaldoxime (IAOx), the common precursor of IGS, camalexin and 4OH-ICN (Mikkelsen et al., 2000), then another cytochrome P450 enzyme, CYP83B1, funnels IAOx into IGS (Naur et al., 2003). To our surprise, no significant increase in expression level of *CYP79B2/B3* was observed, while the transcription level of *CYP83B1* was induced strongly upon infection, and the highest expression occurred at 24 h (Figure 3B). CYP81F2 and CYP81F3, as well as IGMT1/IGMT2, are responsible for side-chain modification of indolic glucosinolate, encoding enzymes involved in the conversion of indole-3-yl-methylglucosinolate (I3G) to 4MI3G. The transcript levels of those genes were intensely induced by *A. brassicicola* infection (Figure 3B).

**Figure 3.**
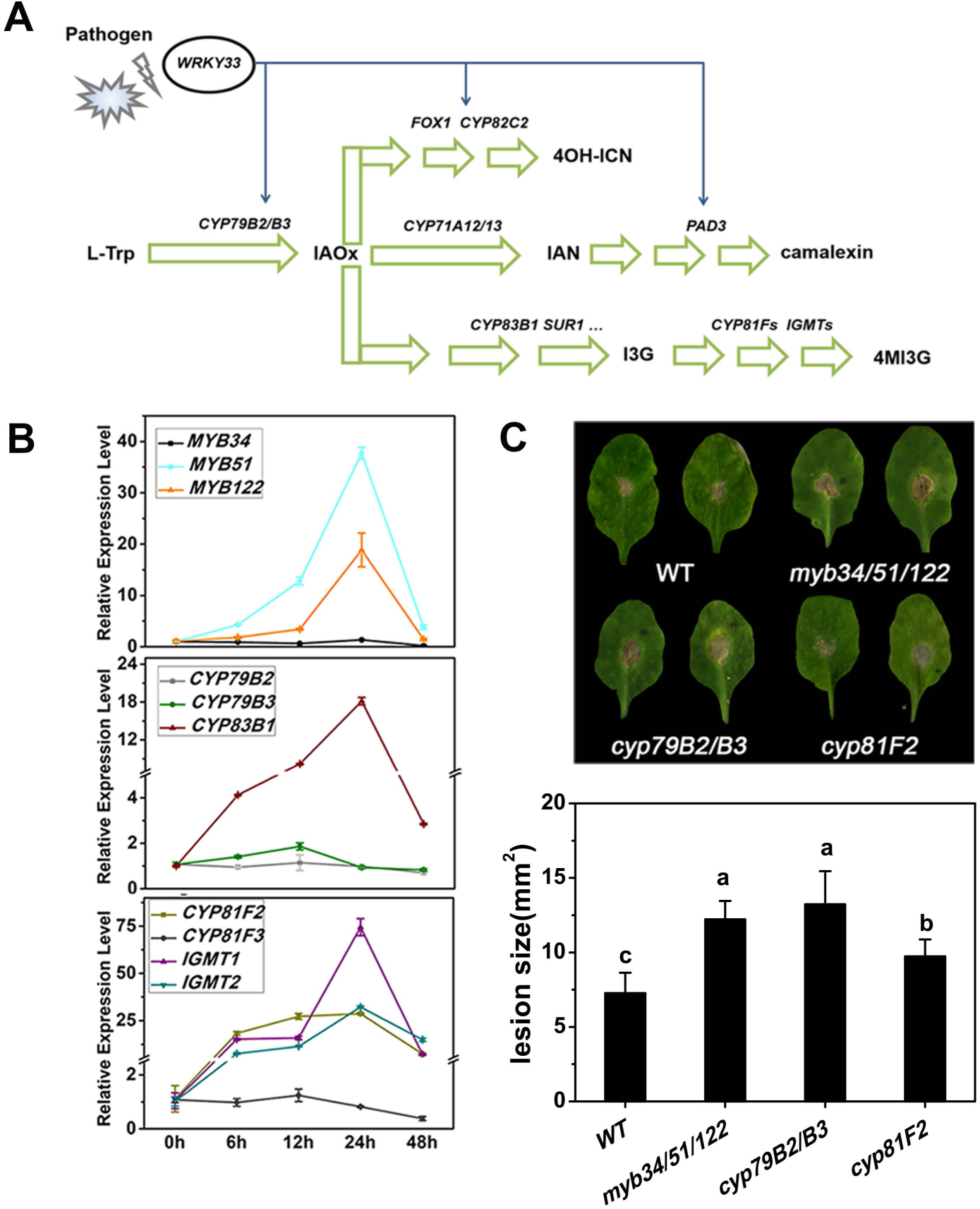
Involvement of tryptophan-derived indolic glucosinolate (IGS) in host defense against *Alternaria brassicicola*. A: Simplified scheme of WRKY33-regulated Trp-derived metabolic pathways. B: qRT-PCR analysis of *A. brassicicola*-induced expression of IGS regulatory and biosynthetic genes in 10 days old WT *Arabidopsis* at indicated time points post fungal spore application. C: Typical phenotypes of 4 weeks old WT, *myb34/51/122, cyp79B2/B3* and *cyp81F2* mutant plants. Plants were infiltrated with *A. brassicicola* in leaves, and then photographed at 5 days post-injection (dpi). Different letters indicate statistically significant differences (P <0.05).

The strong induction of IGS biosynthetic genes in response to *A*. *brassicicola* infection motivated us to assess the contribution of IGS metabolism pathway in host defense against *A*. *brassicicola.* Disease symptom assay was conducted on *myb34/51/122, cyp79B2/B3* and *cyp81F2* (Figure 3C). Lesion sizes in leaves of *myb34/51/122* and *cyp79B2/B3* were significantly larger than in those of WT. Since MYB34, MYB51 and MYB122 not only modulate biosynthesis of indolic glucosinolate but also other Trp-derived metabolites (eg. camalexin and 4OH-ICN), and CYP79B2/B3 regulate biosynthesis of the common precursor (IAO_X_), we included *cyp81F2* which was only deficient in 4MI3G in disease symptom assay. Notably, lesions in *cyp81F2* leaves were larger than that in WT but smaller than that in *myb34/51/122* and *cyp79B2/B3*. This result further confirmed that specified IGS pathway plays an important role in host resistance to *A*. *brassicicola.* Taken together, Trp-derived secondary metabolites, especially 4MI3G as an indolic glucosinolate, contribute to host resistance to *A*. *brassicicola* in *Arabidopsis*.

### PEN2-mediated indolic glucosinolate metabolism is essential for host defense against *Alternaria brassicicola*

To gain an insight into the specific contribution of IGS in resistance to *A. brassicicola*, we measured the levels of indole-3-yl-methylglucosinolate (I3G), 4-methoxyindole-3-yl-methylglucosinolate (4MI3G), and 1-methoxyindol-3-ylmethylglucosinolate (1MI3G), as well as total IGS upon infection at different time points. The steady-state levels of 4MI3G were detected, while 1MI3G, I3G and total IGS levels were gradually decreased (Figure 4A). These results were inconsistent with the significant increase in transcription levels of genes involved in IGS biosynthesis in response to *A. brassicicola* inoculation (Figure 3A), which prompted us to further investigate if IGS hydrolysis functions in resistance to *A. brassicicola*.

**Figure 4.**
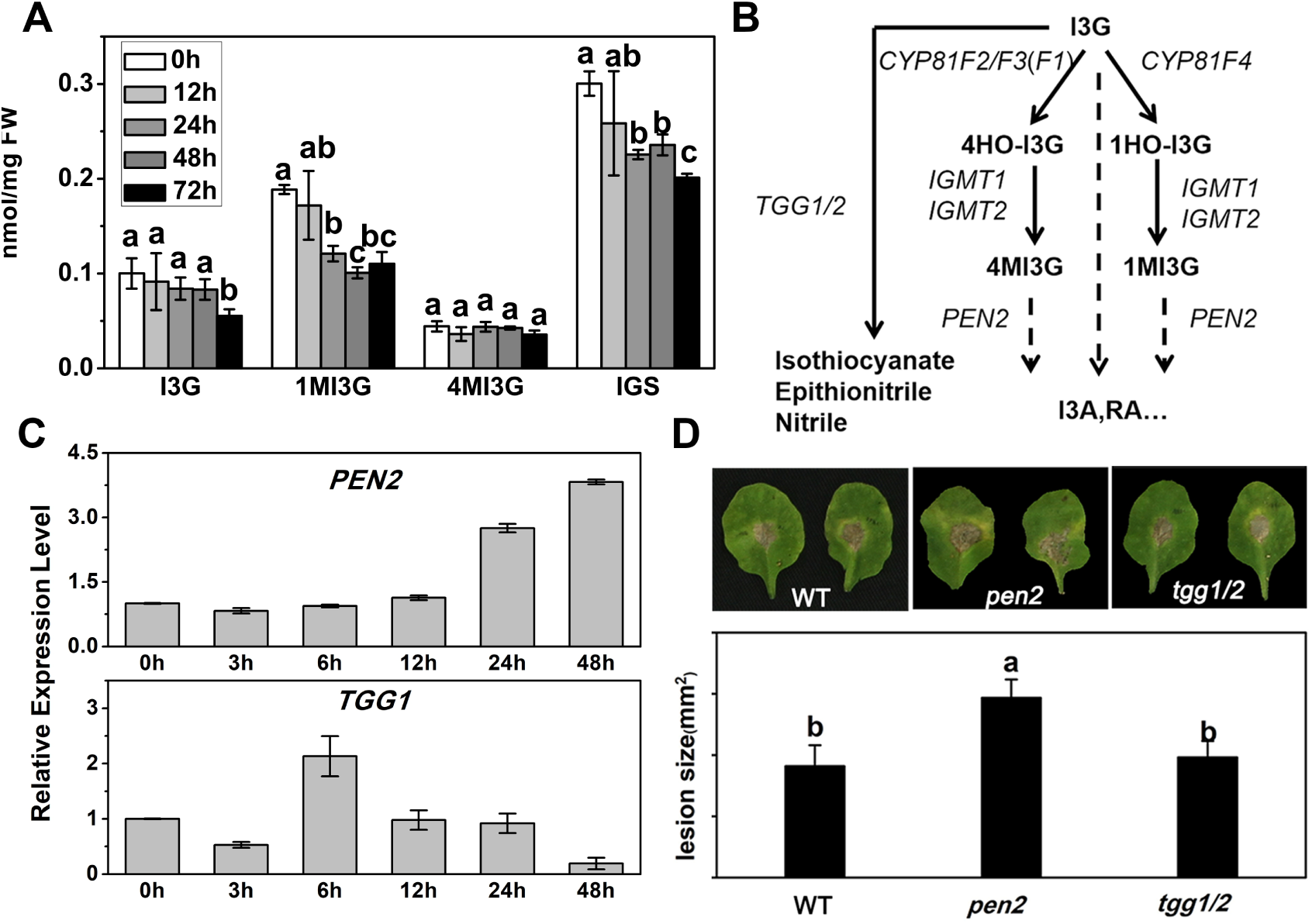
PEN2-mediated indolic glucosinolate (IGS) metabolism is important in host defense against *Alternaria brassicicola*. **A:** Contents of individual and total IGS in 10 days old WT seedlings infiltration with *A. brassicicola* at indicated time points post fungal spore application. Data represent mean ± SE of 4 biological replicates. **B:** Simplified scheme of IGS hydrolysis pathway and related genes in *Arabidopsis*. **C:** qRT-PCR analysis of *A. brassicicola*-induced expression of *PEN2* and *TGG1* in 10 days old WT at indicated time points post fungal spore application. (d) The disease symptoms of 4 weeks old WT, *tgg1/2* and *pen2* mutant plants inoculated with *A. brassicicola* for 5 days. Different letters indicate statistically significant difference (P <0.05)

PENETRATION 2 (PEN2), a thioglucosidase with β-glucosidase activity which was called atypical myrosinase, has been implicated in *Arabidopsis* immunity against pathogens (Lipka et al., 2005). TGGs encode typical myrosinases which are mainly involved in glucosinolate hydrolysis upon tissue damage (Barth and Jander, 2006) (Figure 4B). Thus, we first measured the transcript levels of *PEN2* and *TGG1* in 10-day-old WT upon inoculation with *A. brassicicola*. *PEN2* and *TGG1* were both induced significantly during infection (Figure 4C). These results indicated that both of them might contribute to host resistance to *A. brassicicola.* We then performed disease symptom assay using *pen2* and *tgg1/2* mutants that are impaired in atypical and typical glucosinolates hydrolysis, respectively. Lesion sizes in *pen2* were significantly larger than that in WT, while no significant difference was observed between *tgg1/2* and WT (Figure 4D). Above results suggest that *PEN2*-dependent atypical hydrolysis of IGS is crucial for host defense against *A. brassicicola* in *Arabidopsis*.

### Enhanced biosynthesis of 4MI3G by WRKY33 improves host resistance to *Alternaria brassicicola* both in *Arabidopsis* and Chinese kale

The induction of *WRKY33* expression peaked earlier than those of IGS pathway genes inspired us to explore the relationship between them (Figure 2A and Figure 3B). To gain insight into the role of WRKY33 in IGS biosynthesis, the composition and contents of individual and total IGS, as well as the transcript levels of IGS pathway genes, were analyzed in infected WT and *wrky33*. The contents of individual and total IGS were significantly reduced by the disruption of *WRKY33*. The I3G levels were significantly lower in *wrky33* than in WT upon *A. brassicicola* infection at different time points. Lower levels of 4MI3G in *wrky33* before 24 h compared to the WT were also observed (Figure 5A). The expression levels of IGS pathway genes were highly induced by *A. brassicicola* infection in WT and *wrky33* (Figure 5B). There was a 3.8-fold increase in *MYB51* expression in WT, but only a 2.2-fold increase in *wrky33* at 24 h. A similar expression pattern was also observed in *CYP81F2*, *IGMT1, IGMT2* and *PEN2.* In addition, the expression levels of *CYP83B1,* instead of *CYP79B2*/*B3* was much lower in *wrky33* than that in WT with *A. brassicicola* infection, indicating that WRKY33 serves as a condition-dependent master regulator for the metabolic flux from IAOx to IGS in response to *A. brassicicola*. To verify if WRKY33 has a conserved function in Chinese kale, we performed qRT-PCR analysis in different *BoWRKY33*-silenced Chinese kale lines (pTY-BoWRKY33-2 and pTY-BoWRKY33-3) and the control after inoculation with *A. brassicicola.* The relative expression levels of IGS biosynthetic and hydrolytic genes, namely, *BoMYB51.1, BoCYP81F2, BoIGMT1, BoIGMT2*, and *BoPEN2*, were all significantly reduced in *BoWRKY33*-silenced Chinese kale lines (Figure 5C).

**Figure 5.**
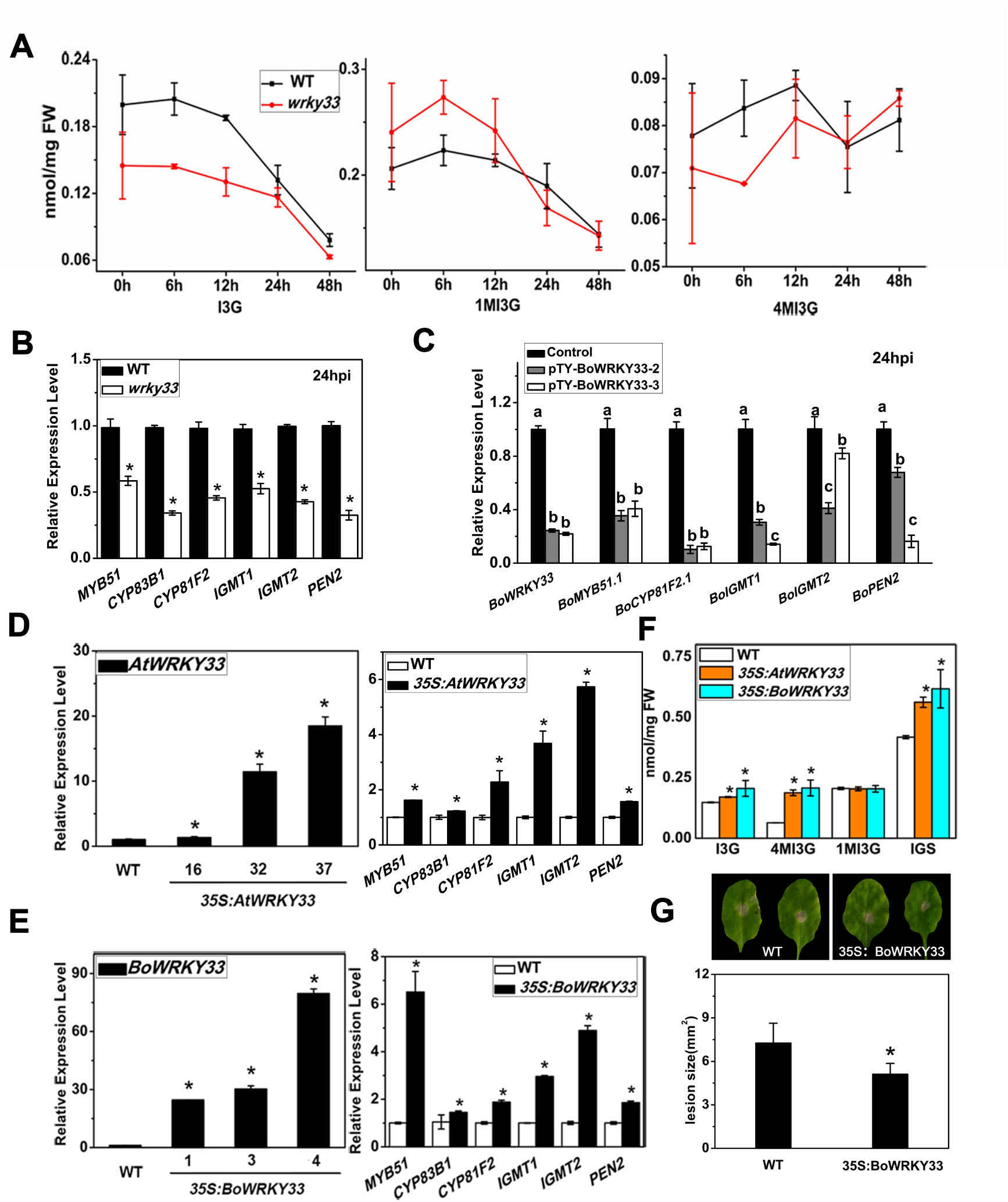
WRKY33 plays an important role in the biosynthesis of 4MI3G to improve host resistance to *Alternaria brassicicola*. **A, B:** IGS contents (A) and qRT-PCR analysis of IGS metabolic genes (B) in *Arabidopsis* WT and *wrky33* at 24h post inoculation with *A. brassicicola*. **C:** qRT-PCR analysis of IGS metabolic genes in control and *BoWRKY33*-silenced Chinese kale lines. **D, E:** qRT-PCR analysis of *AtWRKY33, BoWRKY33*, and IGS metabolic genes in *35S:AtWRKY33* (D) and *35S:BoWRKY33* lines (E) respectively. Bars with asterisk indicate statistically significant differences (P <0.05). **F:** IGS contents in representative *35S:AtWRKY33* and *35S:BoWRKY33* lines. **G:** Disease symptoms and lesion size of 4 weeks old WT and *35S:BoWRKY33 Arabidopsis* at 5 days after inoculation with *A. brassicicola.* Data for the *35S:BoWRKY33 Arabidopsis* are representative of three independent transgenic lines.

In contrast to the results shown in *wrky33* mutants, IGS contents and transcript levels of IGS metabolic genes increased in *AtWRKY33* overexpressing (*35S:AtWRKY33*) lines (Figure 5D). Next, we expressed *BoWRKY33* heterologously in *Arabidopsis* WT, and similar results were observed with *35S:AtWRKY33* lines (Figure 5E). We then analyzed IGS composition and content in *35S:AtWRKY33* and *35S:BoWRKY33* lines, and found that the contents of 4MI3G and total IGS were highly increased in comparison with that in WT (Figure 5F). Furthermore, *BoWRKY33* overexpression *Arabidopsis* plants were more resistant to *A. brassicicola* than the wild type based on disease symptoms and lesion sizes (Figure 5G). These results again indicated that WRKY33 is a positive regulator of IGS de novo biosynthesis and side chain modification to 4MI3G in response to *A. brassicicola* both in *Arabidopsis* and Chinese kale.

### WRKY33 directly binds the promoter of indolic glucosinolate metabolic genes in vivo and in vitro

WRKY TFs specifically bind to motif-containing W-box core sequences, and WRKY33 has been reported to bind to W-box containing the promoter region of *MYB51*, *CYP79B2*, and *CYP79B3* in response to different pathogens (Barco and Clay, 2020). MYB51 is the common TF of Tryptophan-derived secondary metabolites. Our ChIP-qPCR assay, transient expression assay in *N. benthamiana* and yeast one-hybrid assay verified that WRKY33 could directly bind the promoter of *MYB51* (Figure 6A, 6B and 6C). To assess if WRKY33 could directly regulate IGS-related genes induced by *A. brassicicola*, we conducted ChIP-qPCR analysis in WT and *35S:AtWRKY33* lines. We found W-box containing regions within 2000 bp upstream of the transcriptional start site of *CYP83B1*, *CYP81F2*, *IGMT1* and *IGMT2* genes (Figure 7A). The stronger enrichment at W-box containing regions of these genes was observed in *35S:AtWRKY33* in comparison with the WT, indicating that they were direct targets of WRKY33 (Figure 7B). We then verified the effect of WRKY33 on transcriptional function of *CYP83B1*, *CYP81F2*, *IGMT1* and *IGMT2* by using the transient expression assay in *N. benthamiana.* We generated *CYP83B1_pro_:LUC, CYP81F2_pro_ :LUC, IGMT1_pro_ :LUC* and *IGMT2_pro_ :LUC* in which luciferase (LUC) was fused with the promoter. Coexpression of *CYP83B1_pro_ :LUC, CYP81F2_pro_ :LUC, IGMT1_pro_ :LUC* and *IGMT2_pro_ :LUC* with WRKY33 led to an obvious increment of luminescence intensity compared to empty construct control (Figure 7C and 7D), suggesting that the induced expression of *CYP83B1_pro_ :LUC, CYP81F2_pro_ :LUC, IGMT1_pro_ :LUC* and *IGMT2_pro_ :LUC* by WRKY33 is necessary for resistance to pathogens and indole glucosinolate biosynthesis (Bednarek et al., 2009; Clay et al., 2009). The ChIP-qPCR, transient expression assay in *N. benthamiana* and yeast one-hybrid assay were also conducted to test the direct transcriptional regulation of *PEN2* by WRKY33 (Figure 8A, 8B and 8C). Moreover, the ChIP-qPCR analysis was conducted in *35S:BoWRKY33* and WT to test the binding of BoWRKY33 with the promoters of IGS pathway genes. As shown in Supplemental Figure 3, BoWRKY33 also strongly bound to the W-Box containing regions of promoter in IGS pathway genes (*MYB51*, *CYP83B1*, *CYP81F2, IGMT1, IGMT2*, and *PEN2*). These results suggest that WRKY33 is a positive regulator of IGS metabolism both in *Arabidopsis* and Chinese kale (Figure 9)

**Figure 6.**
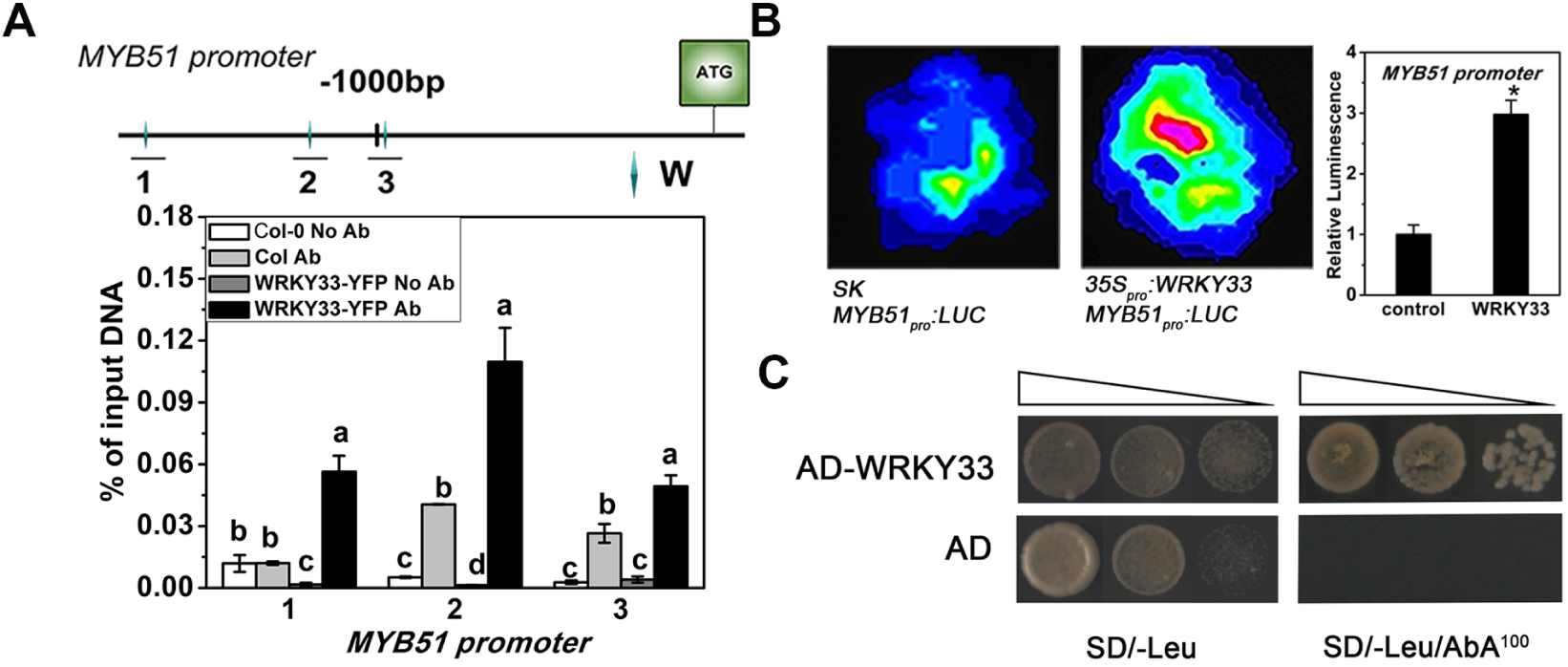
WRKY33 directly binds the promoter of *MYB51*. **A:** Schematic of *MYB51* promoter. The small rhombuses represent W-box motifs in the schematic diagram of the promoters. ChIP assay was performed on WT and *35S:AtWRKY33*. **B:** Transient expression assays showing that WRKY33 actives *MYB51* promoter expression. Representative images of *N. benthamiana* leaves 72 h after infiltration are shown. **C:** Y1H assay showing the binding of WRKY33 to the MYB51 promoter. *AD*, Empty vector used as the control; *AD-WRKY33*, prey vector containing *WRKY33*; SD/-Leu/AbA^100^, we used SD medium without Leu supplemented with 100 ng mL^-1^AbA.

**Figure 7.**
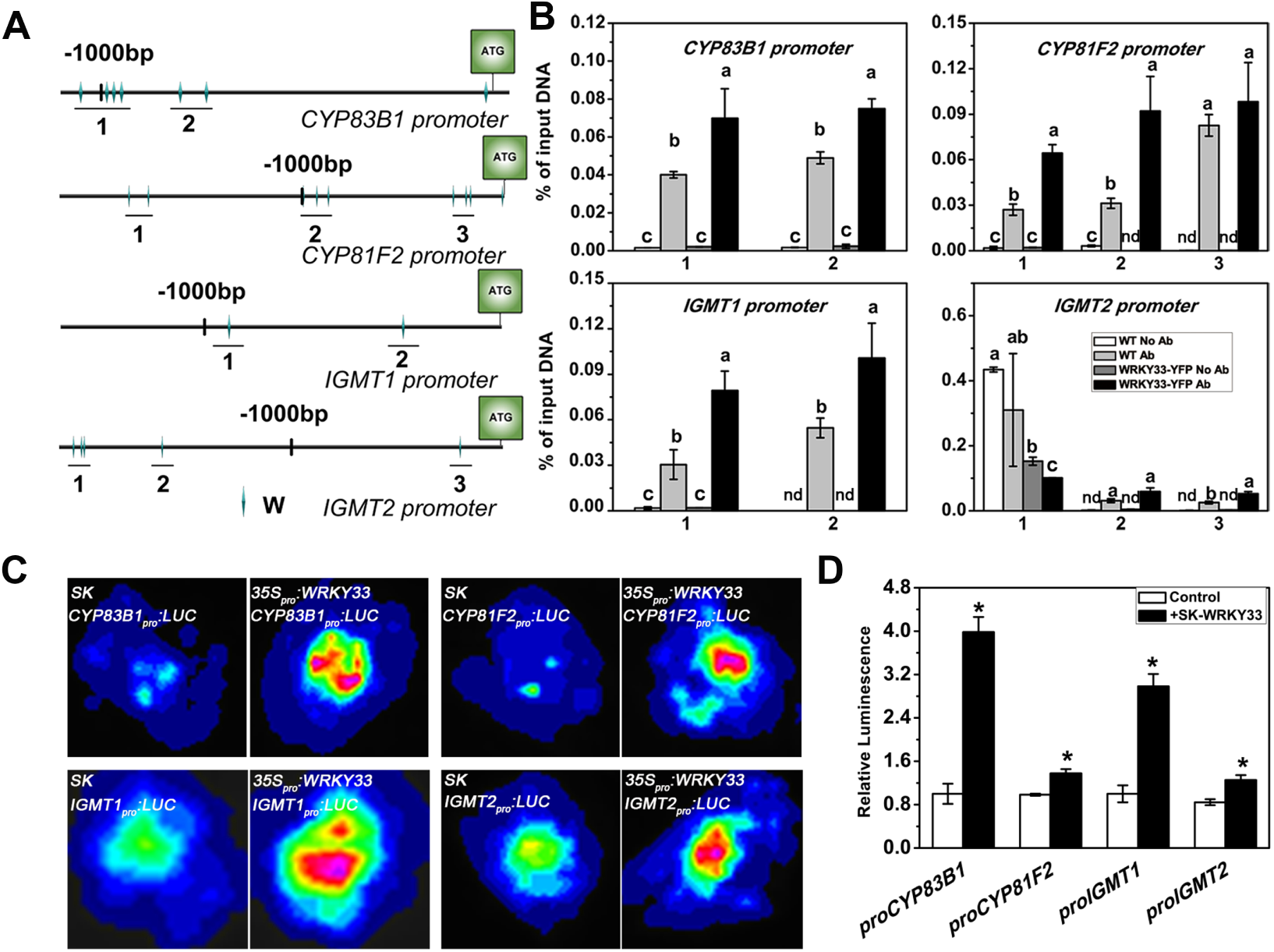
WRKY33 directly binds the promoter of indolic glucosinolate (IGS) metabolic genes in vivo and in vitro. **A:** Schematic of *CYP83B1, CYP81F2, IGMT1*, and *IGMT2* promoters. The small rhombuses represent W-box motifs in the schematic diagram of the promoters. **B:** ChIP assay was performed on WT and *35S:AtWRKY33.* (c-d) Transient expression assays showing that WRKY33 actives *CYP83B1, CYP81F2, IGMT1, and IGMT2* promoters expression.

**Figure 8.**
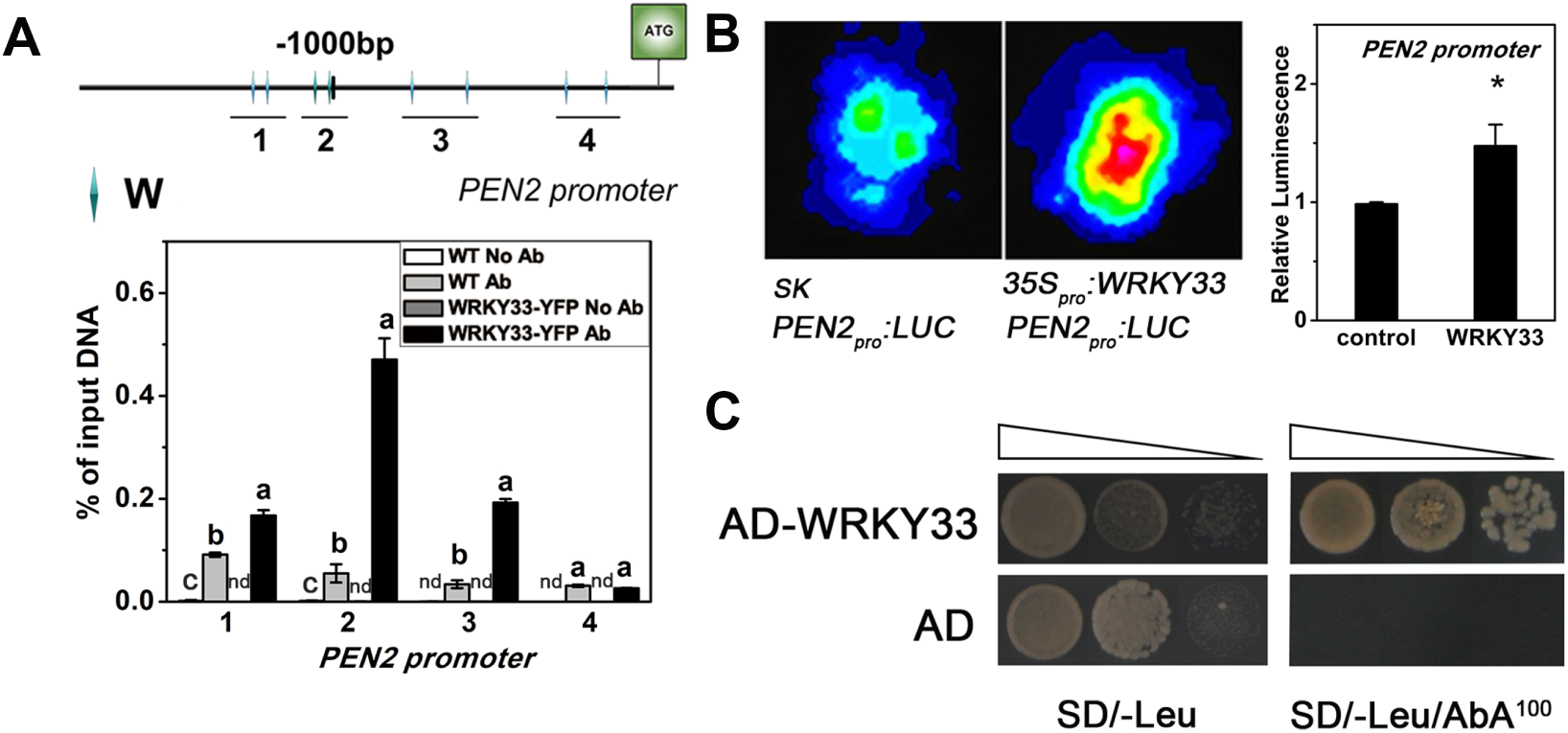
WRKY33 directly regulates transcription of *PEN2*. **A:** Schematic of *PEN2* promoter. The small rhombuses represent W-box motifs in the schematic diagram of the promoters. ChIP assay was performed on 10 days old WT and *35S:AtWRKY33.* Gene expression was quantified by qRT-PCR and calculated as percentages of the input DNA. *P < 0.05. **B:** Transient expression assays showing that WRKY33 actives *PEN2* promoter expression. Representative images of *N. benthamiana* leaves 72 h after infiltration are shown. **C:** Y1H assay have shown the binding of WRKY33 to the *PEN2* promoter. *AD*, Empty vector used as the control; *AD-WRKY33*, prey vector containing *WRKY33*; SD/-Leu/AbA^100^, we used SD medium without Leu supplemented with 100 ng mL^-1^ AbA.

**Figure 9.**
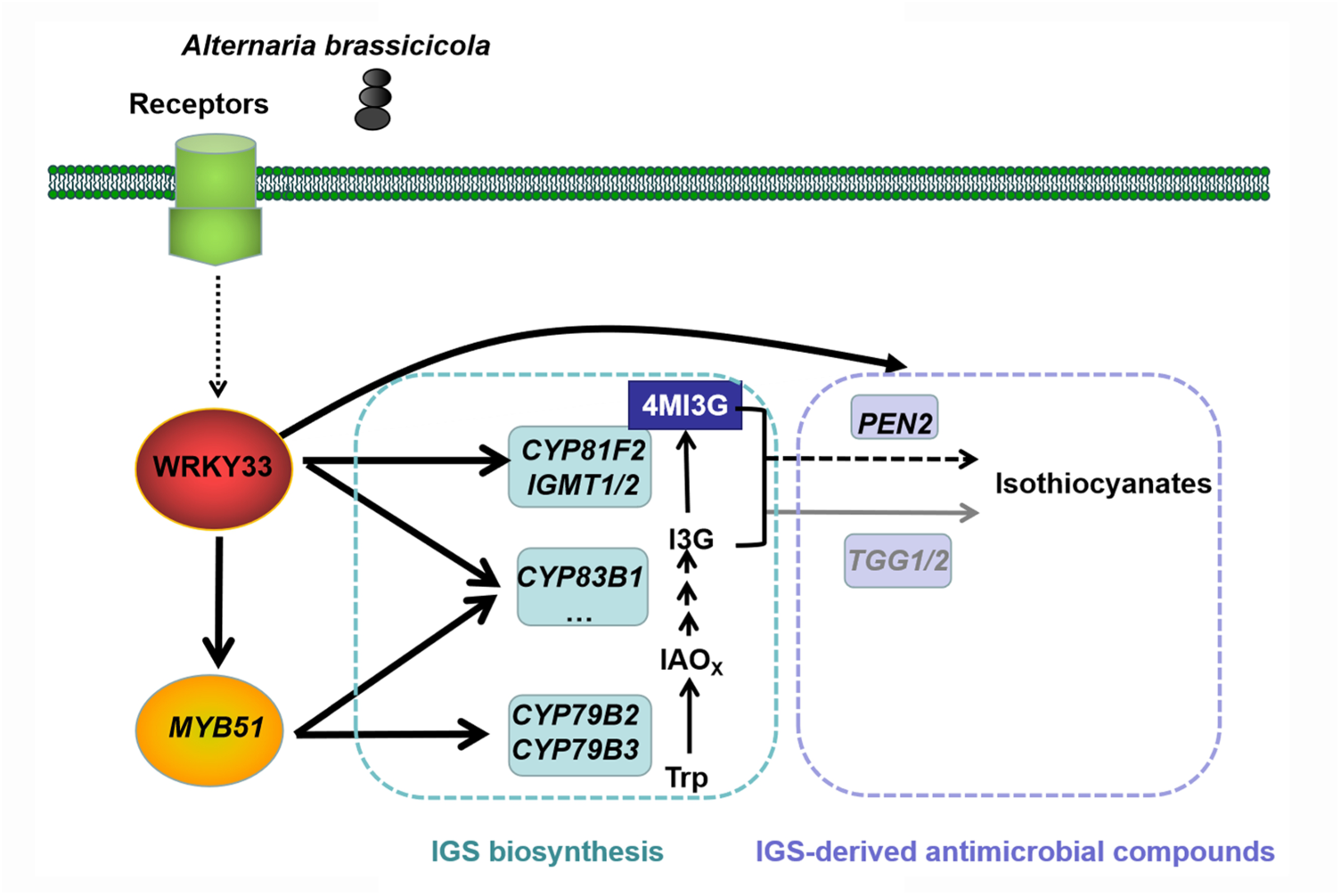
A model depicting the function of WRKY33-regulated indolic glucosinolate (IGS) metabolism in response to *Alternaria brassicicola*. In response to *A. brassicicola*, WRKY33 actively induces the expression of *MYB51* and *CYP83B1* which promotes the biosynthesis of indole-3-yl-methylglucosinolate (I3G), the precursor of, 4-methoxyindole-3-yl-methylglucosinolate (4MI3G). In the meantime, WRKY33 also activates the expression of *CYP81F2*, *IGMT1*, and *IGMT2* to drive side chain modification, converting I3G to 4MI3G. After hydrolyzing by *PEN2* which is also induced by WRKY33 at the level of transcription, IGS are degraded to unidentified unstable compounds to inhibit the pathogen growth.

### The loss of WRKY33-regulated camalexin and 4OH-ICN attenuates host resistance to *Alternaria brassicicola* in Brassica crops

WRKY33-regulated indole glucosinolate metabolic pathway in response to *A. brassicicola* was similar between *Arabidopsis* and Chinese kale. To further investigate which causes the different disease symptoms upon *A. brassicicola* infection in these two Brassica plants (Figure 1A), we first check the protein sequence of WRKY33.

The members of WRKY gene family are characterized by the presence of the highly conserved WRKY domain, which contains the hallmark heptapeptide WRKYGQK and a zinc-finger motif. We compared the structure of WRKY33 in selected Brassica plants through amino acid alignment, and found that they all contain two WRKY domains with highly conserved amino acid sequences, indicating the conservation of WRKY33 among different Brassica species (Supplemental Figure 4). We also analyzed IGS-related genes in Brassica plants. As shown in Supplemental Figure 5, one or more homologous genes could be found in *Brassica oleracea*. For example, *AtMYB51* has two homologous genes *BoMYB51.1* (Bol036286) and *MYB51.2* (Bol030761). *CYP79B2*, *CYP81Fs* and *IGMT5* also have two homologous genes. Other IGS biosynthetic genes were all retained in *Brassica oleracea*. The fact that WRKY33 and IGS are conserved during evolution suggest the possibility that additional factors are involved in difference in host resistance to *A. brassicicola* among difference Brassica plants.

Besides IGS, camalexin and 4OH-ICN are also Trp-derived defense-related metabolites (Bednarek et al., 2009; Clay et al., 2009; Rajniak et al., 2015; Pastorczyk et al., 2019). To explore the phylogenetic distribution pattern of camalexin biosynthesis, we profiled camalexin metabolites in close and distant relatives of *A. thaliana* in response to *A. brassicicola* (Figure 10A). Camalexin biosynthesis was observed across multiple close relatives of *A. thaliana*, while it cannot be detected in Brassica crops. In *A. thaliana*, *PAD3* resides in a near-tandem cluster with paralog *CYP71B28*, which is a putative cytochrome P450 enzyme encoding protein with 77 % identity to *PAD3*. Although *CYP71B28* is similar to *PAD3* in sequence, it is not functionally redundant with *PAD3.* We performed phylogenetic analyses to identify putative *PAD3* orthologs in several relatives of *A. thaliana*. *PAD3* is not present in *Brassica oleracea, Brassica rapa,* and *Brassica napus* (Figure 10B). To be noted, *CYP82C2* has syntenic orthologs only within the *Arabidopsis* genus during the evolution of Brassica plants, leading to undetectable of 4OH-ICN in Brassica crops (Barco et al., 2019). Thus, only the genes related to glucosinolate metabolic pathway, but not the camalexin and 4-OH-ICN pathway, retained and duplicated during the polyploid genome evolution of Brassica species.

Our HPLC analysis of camalexin and IGS in *Arabidopsis* and Chinese kale also showed that camalexin was only detected in infected *Arabidopsis,* while IGS was detected in both *Arabidopsis* and Chinese kale after infection (Supplemental Figure 6). These results again highlight the crucial role of IGS metabolic pathway in Brassica crops. Because of the loss of Tryptophan-derived camalexin and 4-OH-ICN biosynthesis in Brassica crops (Figure 10C), WRKY33-mediated tryptophan-derived secondary metabolites patterns in response to *A. brassicicola* are simplified. In *Arabidopsis,* WRKY33 was induced to modulate camalexin, 4-OH-ICN and IGS pathway to combat the pathogen upon *A. brassicicola* infection. While in Brassica crops, only WRKY33-regulated IGS pathway contributes to the resistance against *A. brassicicola* (Figure 10D).

## Discussion

Alternaria black spot caused by necrotrophic *Alternaria* pathogens is one of the most serious diseases in economically important Brassica crops. Brassica crops show more severe symptoms upon *A. brassicicola* infection when compared to *Arabidopsis* (Figure 1A). It is of great biological and economic significance to elucidate the mechanism in host resistance to *A. brassicicola* and to understand the causes that lead to the differential susceptability in *Arabidopsis* and Brassica crops. Our study here identified a conserved WRKY33-regulated indolic glucosinolate (IGS) metabolism both in *Arabidopsis* and Brassica crops, while WRKY33-mediated Tryptophan (Trp)-derived camalexin and 4-hydroxy-indole-3-carbonylnitrile (4OH-ICN) are missing in Brassica crops towards *A. brassicicola* (Figure 10).

**Figure 10.**
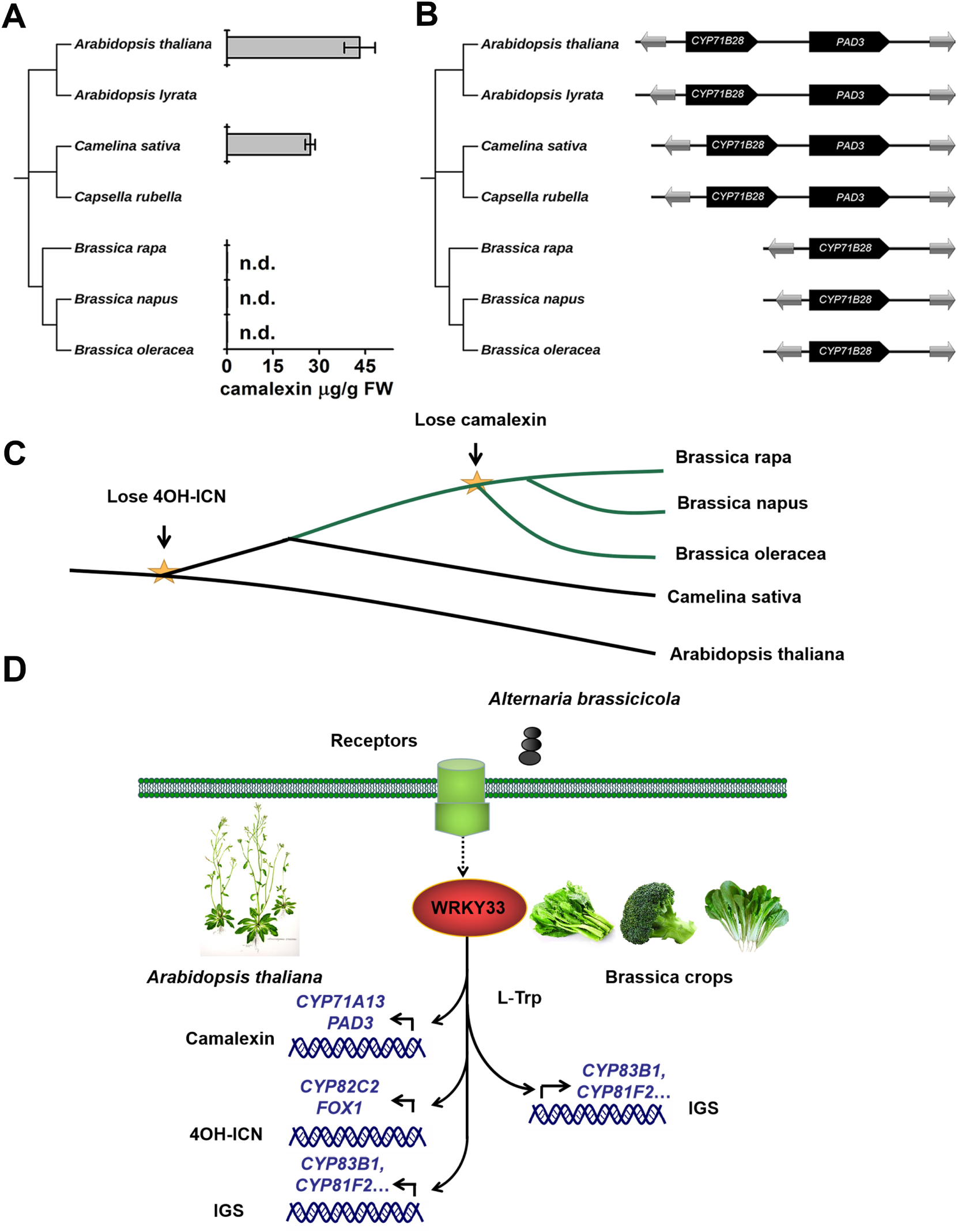
The loss of WRKY33-regulated camalexin and 4OH-ICN attenuates the resistance of Brassica crops to *Alternaria brassicicola*. **A:** Phylogenetic species tree and HPLC analysis of camalexin in selected Brassica plants inoculated with *A. brassicicola* for 48 h, not detect (n.d.). **B:** Synteny map of the *PAD3* and *CYP71B28* genes. C: The phylogenetic relationship of Brassica crops and closely related Brassica species, the yellow star denotes the evolutionary node of the Tryptophan-derived secondary metabolites. D: Different patterns of WRKY33-regulated tryptophan-derived secondary metabolites between *Arabidopsis* and Brassica crops against *A. brassicicola* invasion.

### WRKY33 functions in defense against *Alternaria brassicicola* infection both in *Arabidopsis* and Brassica crops

WRKY transcription factors play vital roles in tuning the cellular responses triggered by pathogens in many plant species (Dong et al., 2003; Chen et al., 2010). WRKY33 is a pathogen-inducible transcription factor, whose expression is regulated by the MPK3/MPK6 cascade (Zhou et al., 2020). WRKY33 was also shown to be essential for the induction of camalexin biosynthesis in *Arabidopsis* infected with *Pseudomonas syringae* and positively regulates host resistance to necrotrophic pathogens *Botrytis cinerea* (Zheng et al., 2006; Lai et al., 2011; Liu et al., 2015). Previous microarray analyses have shown that WRKY33 modulates genes involved in plant hormone pathways and Trp-derived camalexin and 4OH-ICN upon *B. cinerea* or flg22 infection (Birkenbihl et al., 2012; Birkenbihl et al., 2017). Our RNA-seq data showed that WRKY33 and Trp-derived secondary metabolic pathway genes were all induced during fungus infection in *Arabidopsis* (Figure 1). And the symptom assay with *wrky33*+*35S:AtWRKY33* confirmed the positive role of WRKY33 in host resistance (Figure 2A). In Brassica crops, the symptoms of *BoWRKY33-*silenced Chinese kale and *BoWRKY33* overexpression lines further verified the role of BoWRKY33 as a key transcription factor and resistant gene in controlling black spot caused by *A. brassicicola* (Figure 2B and Figure 5G).

### Indolic glucosinolate biosynthesis and PEN2-mediated atypical degradation function in host resistance to *Alternaria brassicicola*

Glucosinolates have been shown to play important roles in plant defense against pathogens (Bednarek et al., 2009). Our RNA-seq data and qRT-PCR analysis indicated that numerous glucosinolates-related genes were induced by *A. brassicicola* infection (Figure 1D and Figure 3B). Previous researches have shown that *CYP79B2* and *CYP79B3* were induced intensively upon *Botrytis cinerea* infection, leading to a significant increase in IAOx, the common precursor of camalexin, 4OH-ICN and IGS (Barco and Clay, 2020). However, *CYP79B2* and *CYP79B3* were only slightly induced, while *CYP83B1,* a gene involved in the specific downstream biosynthetic step from IAOx to IGS, was intensively induced upon *A. brassicicola* infection in the present study (Figure 3B), indicating that IGS pathway might play a more important role in response to host-specific fungi *A. brassicicola,* rather than to wide-ranging host fungi *Botrytis cinerea*. In addition, the *A. brassicicola* infected IGS-related mutants (*myb34/51/122, cyp79B2/B3* and *cyp81F2*) showed enhanced susceptibility to the pathogen (Figure 3C). The disease symptom assays on *Arabidopsis* mutants indicated that Trp-derived secondary metabolites, especially indolic glucosinolates, contribute to host resistance to *A*. *brassicicola* in *Arabidopsis*.

The levels of I3G and 1MI3G were decreased, along with a relatively steady 4MI3G content (Figure 4A), inconsistent with the observation of intensively induced expression of genes involved in IGS biosynthesis post *A. brassicicola* infection (Figure 3B), which makes us further investigate IGS hydrolysis. Our previous report demonstrated that TGG mediated typical hydrolysis of IGS functions in defense against chemical stress induced by mycotoxin fumonisin B1 (Zhao et al., 2015). While our infection assays with *A. brassicicola* showed that *pen2,* the atypical hydrolysis mutant, instead of *tgg1/2* was more susceptible to *A. brassicicola* compared to the WT (Figure 4D), suggesting different mechanisms of IGS metabolism in response to biotic and abiotic stress, respectively.

### Parallel regulation of indolic glucosinolate metabolism by WRKY33 between *Arabidopsis* and Brassica crops

Regulation of tryptophan-derived secondary metabolites is an important approach of WRKY33 to tune plant resistance. It has been reported that 4OH-ICN pathway genes (eg, *CYP82C2*) were recruited to the WRKY33 regulon in *Arabidopsis* in response to *Botrytis cinerea* (Barco et al., 2019). WRKY33 is also a master regulator of camalexin response pathway (Mao et al., 2012) which was activated by MAP kinase signaling upon *Botrytis cinerea* infection, and then directly induced the transcription of camalexin biosynthesis (*CYP71A13* and *PAD3*) and transport (*PEN3* and *PDR12*) genes (Mao et al., 2012; He et al., 2019; Pastorczyk et al., 2019). In addition, WRKY33 can be phosphorylated by CPK5 to promote the expression of biosynthetic genes in camalexin pathway (Zhou et al., 2020). Recent research has shown that MYB51, MYB122, and MYB34 are not required for induction of main genes involved in side-chain modification (*CYP81F2, IGMT1*, and *IGMT2*), suggesting the possible involvement of additional transcription factors in side-chain modification of IGS (Xu et al., 2016). The regulatory role of WRKY33 in biosynthesis (*MYB51* and *CYP83B1*), side-chain modification (*CYP81F2*, *IGMT1* and *IGMT2*) and hydrolysis (*PEN2*) of IGS (Figures 5-8) indicates a hierarchical regulatory of WRKY33 and glucosinolates-related genes in the process. The novel finding that WRKY33 fulfills the regulation of IGS pathway and highlights its central role in modulation of IGS metabolic pathway as well as host resistance to *A. brassicicola*. We also observed a conserved mechanism of BoWRKY33 in controlling biosynthesis and hydrolysis of IGS in Chinese kale which is similar with that in *Arabidopsis thaliana* (Supplemental Figure 3).

As the working model is shown (Figure 9), there is a WRKY33-centraled network in controlling IGS metabolism in response to *A. brassicicola.* Upon *A. brassicicola* infection, WRKY33 induces expression of *MYB51* and *CYP83B1* which promotes the biosynthesis of I3G, the precursor of 4MI3G. WRKY33 also directly activates the expression of *CYP81F2*, *IGMT1*, and *IGMT2* to drive side chain modification of I3G, and produce 4MI3G, which is in turn hydrolyzed by *PEN2*. Then 4MI3G is converted to unidentified unstable compounds which are transported by PEN3 for extracellular defense against fungal pathogen.

### Distinct inducing mechanisms of Trp-derived secondary metabolites by WRKY33 against *Alternaria brassicicola* between *Arabidopsis* and Brassica crops

Camalexin, 4OH-ICN, and IGS can all be detected *in Arabidopsis thaliana*, but in Brassica species such as *Brassica rapa*, *Brassica napus* and *Brassica oleracea*, camalexin and 4OH-ICN are absent upon fungus attack (Sigareva and Earle, 1999; Bednarek et al., 2011; Barco et al., 2019). *CYP82C2* has syntenic orthologs only within the *Arabidopsis* genus during the evolution of Brassica plants, leading to undetectable of 4OH-ICN in Brassica crops (Barco et al., 2019). Our current study also showed that Brassica crops could not synthesize camalexin by the loss of camalexin biosynthetic gene *PAD3* (Figure 10A, 10B and Supplemental Figure 6). Brassica species usually produce brassinin instead of camalexin, as a kind of phytoalexin in response to pathogen (PEDRAS and MINIC, 2012). It has been reported that pathogen-inducible IGS metabolism is evolutionarily ancient in the Brassica plants and has been largely retained among them, whereas camalexin is restricted to the *Camelineae tribe* of *Brassicaceae* (Bednarek et al., 2011). We then analyzed the evolution of transcription factors and biosynthetic genes in IGS pathway in Brassica plants. There are two homologous genes of *MYB51*, *CYP79B2* and *CYP81F2* through tandem duplication in *Brassica oleracea,* and other IGS biosynthetic enzymes are all retained in *Brassica oleracea* (Supplemental Figure 4). Whole-genome duplications, gene rearrangements, and substrate promiscuity potentiated the evolution of glucosinolate biosynthetic enzymes, regulators, and transporters by natural selection and in turn may have led to the repeated evolution of glucosinolate metabolism and diversity in higher plants (Barco and Clay, 2019). Our results are consistent with the former research. Taken together, the biosynthesis of IGS and their derivatives are critical for resistance of Brassica crops.

WRKY33 initiates a distinct regulation of Trp-derived secondary metabolites between *Arabidopsis* and Brassica crops against *A. brassicicola* invasion. The regulation of WRKY33 on camalexin and 4OH-ICN biosynthesis is consistent with previous researches in *B. cinerea* or flg22-elicited *Arabidopsis* plants (Mao et al., 2012; Barco et al., 2019). Our data also showed the massive induction of camalexin and 4OH-ICN biosynthetic genes in *Arabidopsis* after *A. brassicicola* infection and the fine regulation of WRKY33 on IGS (Figure 1). Thus, WRKY33-regulated camalexin, 4OH-ICN and IGS pathways all participate in host resistance to the invasion of *A. brassicicola* in *Arabidopsis*, but only WRKY33-regulated IGS pathway contributes to host resistance in Brassica crops (Figure 10). It highlights the importance of WRKY33-regulated IGS metabolic pathway in host resistance of Brassica crops to *A. brassicicola*. The severer symptom in Brassica crops probably due to the loss of camalexin and 4-OH-ICN biosynthetic genes *PAD3* and *CYP82C2* during the process of domestication and evolution. They are potential targets in developing a green and sustainable disease resistance against *A. brassicicola* in Brassica crops. Our results provide a strategy for enhancing the resistance of Brassica crops to black spot disease by improvement of WRKY33-mediated Trp-derived secondary metabolic pathways. Future insights into the transgenic Brassica crops will be critical to the evolution of pathogen-inducible tryptophan metabolic pathways and their functions in plant innate immunity.

## Conclusion

In summary, we revealed that WRKY33 directly regulates indolic glucosinolate (IGS) biosynthesis and atypical hydrolysis, conferring to host resistance to *Alternaria brassicicola* both in *Arabidopsis* and Chinese kale (Figure 9). We also observed a severer symptom in Chinese kale (*Brassica oleracea* var. *alboglabra* Bailey) than in *Arabidopsis.* Comparative analyses of the origin and evolution of Trp-metabolism in different Cruciferous plants further elucidated a novel WRKY33-mediated distinct immune system between *Arabidopsis* and Brassica crops against *Alternaria brassicicola*, via induction of three above Trp-derived secondary metabolites simultaneously in *Arabidopsis*, while only IGS in Brassica crops due to evolution and domestication (Figure 10). Our results highlight the significance of glucosinolate metabolic regulatory network in plant immunity, and are potential in developing a green and sustainable disease control strategy towards *Alternaria brassicicola* in Brassica crops.

## Materials and Methods

### Plant materials and growth conditions

Seeds were sterilized for 12 min in 10% bleach and washed with sterile water 6 times. The seeds were stratified for 3 d at 4°C and planted on ½ MS (Murashige Skoog) growth medium. For qRCR and glucosinolate assays, *Arabidopsis* plants were grown on ½ MS medium under a photoperiod of 16 h light/8 h dark for 10 days in a plant growth chamber at 21°C. For disease assays, plants were grown on soil under a 12 h light/12 h dark light cycle. Chinese kale (*Brassica oleracea* var. *alboglabra* Bailey) plants were grown on soil under a photoperiod of 16 h light/8 h dark at 25°C.

All *Arabidopsis thaliana* plants used in this study are in the Columbia (Col-0) background. Seeds of *wrky33-2* (GABI_324B11) were kindly provided by Dr. Zhixiang Chen. Mutant seeds of *cyp79B2/B3, myb34/51/122*, *cyp81F2, tgg1/2* and *pen2* were used (Zhao et al., 2015)(Zhao *et al*., 2015).

### Plants transformation

The full length encoding fragments of *AtWRKY33* and the homologous gene in Chinese kale *BoWRKY33* were cloned into pCAMBIA1300-YFP vector to generate *35S:AtWRKY33-YFP* and *35S:BoWRKY33-YFP* fused plasmid for plant transformation. *35S:AtWRKY33-YFP* and *35S:BoWRKY33-YFP* were introduced into Col-0 to generate *35S:AtWRKY33* and *35:BoWRKY33 Arabidopsis. 35S:AtWRKY33-YFP* was introduced into *wrky33* to generate *wrky33*+*35S:AtWRKY33.* The *Agrobacterium tumefaciens* strain GV3101 was used for the transformation experiment, and transformants were selected on agar media containing 15μg/mL hygromycin B (Invitrogen, Carlsbad, CA).

### Virus-Induced Gene Silencing (VIGS) assay

VIGS experiments were performed as described (Jupin, 2013). We used the *Turnip yellow mosaic virus* (TYMV)-derived vector pTY-S to induce VIGS in Chinese kale (*Brassica oleracea* var. *alboglabra* Bailey). The 4-week-old plant leaves were inoculated with empty plasmid pTY-S or plasmid pTY-WRKY33. New leaves were sampled 3 weeks post-inoculation.

### *Alternaria brassicicola* cultivation and disease assays

*Alternaria brassicicola* were cultivated on Potato Dextrose Agar (PDA) and incubated at 25°C. Collection of the *Alternaria brassicicola* spores and plant inoculation were performed as described (Zheng et al., 2006). For disease assays performed on 4-5 weeks-old soil-grown *Arabidopsis* plants and 2 months old Chinese kale, a single 10 μL drop and 100 μL of a suspension of 5×10^5^ spores mL^-1^ in water was placed on each leaf respectively. After 5 days, lesions were photographed and lesion sizes were measured using ImageJ. For qRT-PCR assays and determination of glucosinolates, 10-day-old *Arabidopsis* seedlings grown on ½ MS growth medium were transferred into flasks with 2 mL fungal solution with a density of 5×10^5^ spores mL^-1^ and cultured for 24hpi with gently shaking.

### RNA-seq assay

Total RNA was extracted from mock-treated (water for 24h) and *Alternaria brassicicola* (5×10^5^ spores mL^-1^ for 24 h) infected 10-day old plants (Col-0 and *wrky33*). Total reads were mapped to the *Arabidopsis* genome (TAIR10). The sequencing library was then sequenced on a Hiseq X platform (Illumina) by Shanghai Personal Biotechnology Cp. Ltd.

### RNA extraction and qRT-PCR

Total RNA was extracted from 10-day-old seedlings with RNAiso Plus (Takara, Japan) according to the manufacturer’s instruction. RNA samples were reverse-transcribed into cDNA using Prime Script RT Master Mix (Takara). The expression level of *Arabidopsis ACTIN7* or Chinese kale *BoACTIN7* were used as internal control and the expression levels of other genes were computed with the 2^−ΔΔCT^ method. The Primers used in the current study are listed in Table. S2.

### Glucosinolates assay

Glucosinolates were extracted and analyzed as previously described with some modifications (Zhao et al., 2015). Samples (0.1g) were homogenized in 1 mL of 90% methanol at 60Hz for 1min. And then another 1 mL of 90% methanol was added. The extracts were incubated at room temperature for 1 hour. After centrifugation (1 min, 10000 g), the supernatant was transferred into a new 2 mL tube. 1 mL of the combined extracts was applied to a DEAE-Sephadex A-25 column. The column was washed with 1 mL of 90% methanol and then with 1mL of water. After washing, 100 μL of 0.1% (1.4units) aryl sulphatasethe were added into the column and incubated for 12-14 h to glucosinolates to their desulpho analogs. The desulpho analogs were collected by washing the column with 1 mL of water. The high-performance liquid chromatography (HPLC) analysis was carried out by using an HPLC instrument (Shimadzu, Kyoto, Japan) with SPD-M20A diode array detector. Data were given as nmol/mg fresh weight (FW).

### Camalexin analysis

Camalexin was extracted and analyzed as previously described with some modifications (Yang et al., 2020). After treatment, the samples were diluted with methanol and analyses on HPLC instrument (Shimadzu, Kyoto, Japan) with SPD-M20A diode array detector. Data were given as μg/g fresh weight (FW).

### ChIP-qPCR assay

ChIP experiments were performed as described previously with some modifications (Zhai et al., 2015). 2 grams of 10-day-old *35S:AtWRKY33, 35S:BoWRKY33* and the wild-type Col-0 seedlings were used for ChIP assay. YFP antibody (Abcam) was used to immunoprecipitate the protein-DNA complex. Chromatin precipitated without antibody was used as the negative control, while the isolated chromatin before precipitation was used as input control. The enrichment was calculated as percentage of the input control. Primers used for ChIP-qPCR were listed in Table. S3.

### Transient expression assays in *Nicotiana benthamiana* leaves

The transient expression assays were d in *N. benthamiana* leaves as previously described with minor modifications (Zhai et al., 2015). The promoters of *MYB51, CYP83B1, CYP81F2, IGMT1, IGMT2,* and *PEN2* were amplified and cloned into the vector pGreenII 0800-LUC to generate the reporter constructs *MYB51pro:LUC*, *CYP83B1pro:LUC*, *CYP81F2pro:LUC*, *IGMT1pro:LUC, IGMT2pro:LUC and PEN2pro:LUC.* The coding sequence of *WRKY33* was PCR-amplified and cloned into pGreenII 0029-62-SK to generate the effector construct. We used a low-light cooled CCD imaging apparatus (NightOWL II LB983) to get the LUC image and to count luminescence intensity. Each tobacco leave was sprayed with 100 mM luciferin (Promega) and placed in darkness for 5 min before luminescence detection.

### Yeast one-hybrid assays

Yeast one-hybrid assay was performed as the Clontech Yeast One-Hybrid System manufacturer’s protocol. The promoter fragments of *MYB51 and PEN2* were cloned into pAbAi vector, and the coding sequence of WRKY33 was amplified cloned into pGADT7 vector to generate a prey construct. The empty vector (AD) was served as the negative control. We used SD medium without Leu supplemented with 100 ng mL^-1^ AbA, transformed yeast cells were dotted at 1, 10^-1^, and 100^-1^ dilutions on the SD/-Leu/AbA^100^ medium.

### Statistical analysis

Statistical analysis was performed using the SPSS package program version 11.5 (SPSS Inc., Chicago, IL, USA). The differences between two groups of data at a specific time point were analyzed by using Student’s *t* tests. Single asterisks above the columns indicate differences that are statistically significant (*P < 0.05). For more than two samples were compared, differences were determined by one-way analysis of variance (ANOVA), followed by the least significant difference (LSD) test at a 95% confidence level (*P* < 0.05). Different letters above the columns are used to indicate statistically significant differences. The values are reported as means with SE for all results.

### Accession Numbers

WRKY33 (AT2G38470), MYB34 (AT5G60890), MYB51 (AT1G18570), MYB122 (AT1G74080), MYB28 (AT5G61420), MYB29(AT5G07690), MYB76 (AT5G07690), CYP79B2 (AT4G39950), CYP79B3 (AT2G22330), CYP83B1 (AT4G31500), CYP81F2 (AT5G57220), CYP81F3 (AT4G37400), CYP81F4 (AT4G37410), IGMT1(AT1G21100), IGMT2 (AT1G21120), PEN2 (AT2G44490), TGG1 (AT5G26000).

## Supplemental Data

**Supplemental Figure S1.** GO enrichment of wrky33 VS WT in response to *Alternaria brassicicola*.

**Supplemental Figure S2.** Camalexin and 4-OH-ICN biosynthetic genes were induced by *Alternaria brassicicola*.

**Supplemental Figure S3.** BoWRKY33 is a transcriptional activator of indolic glucosinolate (IGS) metabolic genes.

**Supplemental Figure S4.** Protein sequence alignment of WRKY33 in selected Brassica plants.

**Supplemental Figure S5.** Phylogenetic history of indolic glucosinolate (IGS)-related genes.

**Supplemental Figure S6.** Camalexin and indolic glucosinolate (IGS) contents of *Arabidopsis* and Chinese kale in response to *Alternaria brassicicola*.

**Supplemental Table S1.** Fold induction of all genes 24 hpi with *Alternaria brassicicola* (p≤0.05).

**Supplemental Table S2.** Primer sequences used in qPCR.

**Supplemental Table S3.** Primer sequences used in ChIP-qPCR.

## Acknowledgments

We thank Prof. Zhixiang Chen (Purdue University, U.S.) for providing *wrky33* (GABI_324B11) and Prof. Mingfang Zhang (Zhengjiang University, China) for providing plasmid pTY-S.

